# ENACT: End-to-End Analysis of Visium High Definition (HD) Data

**DOI:** 10.1101/2024.10.17.618905

**Authors:** Mena Kamel, Yiwen Song, Ana Solbas, Sergio Villordo, Amrut Sarangi, Pavel Senin, Sunaal Mathew, Luis Cano Ayestas, Seqian Wang, Marion Classe, Ziv Bar-Joseph, Albert Pla Planas

## Abstract

**Motivation:** Spatial transcriptomics (ST) enables the study of gene expression within its spatial context in histopathology samples. To date, a limiting factor has been the resolution of sequencing based ST products. The introduction of the Visium High Definition (HD) technology opens the door to cell resolution ST studies. However, challenges remain in the ability to accurately map transcripts to cells and in assigning cell types based on the transcript data.

**Results:** We developed ENACT, the first tissue-agnostic pipeline that integrates advanced cell segmentation with Visium HD transcriptomics data to infer cell types across whole tissue sections. Our pipeline incorporates novel bin-to-cell assignment methods, enhancing the accuracy of single-cell transcript estimates. Validated on diverse synthetic and real datasets, our approach is both scalable and effective offering a robust solution for spatially resolved transcriptomics analysis.

**Availability and implementation:** ENACT source code is available at https://github.com/Sanofi-Public/enact-pipeline. Experimental data is available at https://zenodo.org/records/13887921. Supplementary information: Supplementary data are available at BiorXiv online.

## Introduction

Spatial transcriptomics (ST) is a promising new technology that enables researchers to explore the spatial distribution of cells within tissues. ST technologies can be divided into two categories; sequencing-based (e.g. Visium and GeoMX) and image-based (e.g. Xenium, MERFISH, and CosMX). Sequencing-based technologies provide several advantages including the ability to comprehensively map the entire transcriptome, identification of splice variants, and computation of RNA-velocity (1). The data obtained from sequencing-based methods is collected from spots placed in a grid like manner. These spots typically have a multi-cell resolution, preventing the study of the tissue at a single-cell resolution. Recently, 10X Genomics released Visium HD (2), a new sequencing-based ST platform that collects transcript counts at a sub-cellular resolution. Specifically, spots (also referred to as bins) in Visium HD are 2μm x 2μm. To enable cell-based analysis, Visium HD allows the user to aggregate bins resulting in an 8μm x 8μm dimension for each spot, which roughly covers the size of a cell. However, accurately get-ting single-cell transcript estimates from the aggregated 8μm x 8μm is not trivial. First, some cells are smaller than the aggregated spot size which leads to contamination. Even more problematic, these aggregated spots rarely completely overlap a single cell. In many cases, each such aggregated spot overlaps 2 or more cells and vice versa, each cell overlaps more than 1 spot (often more than 2 given the 3D placement) especially in cases where cells are smaller than 8μm in diameter, or where cells overlap or are tightly spaced.

To address this problem, instead of using the 8μm x 8μm bins for downstream analysis, previous work such as Bin2cell (3) proposes a method that combines morphology and gene expression information to obtain accurate single-cell transcript counts. Bin2cell uses Stardist (4) to obtain cell outlines. These outlines are then expanded, and the 2μm x 2μm bins with centroids within each cell outline are aggregated to obtain single-cell transcript counts.

While such integrated imaging plus sequencing method obtains good results in some cases, as we show it can be less effective when cells are tightly spaced. In such cases many spots overlap multiple cells. Thus, methods for distributing transcripts in such bins between the cells they overlap are required to obtain an accurate representation of the expression of each cell.

To address these issues, we developed ENACT, a comprehensive pipeline for processing Visium HD results. Our pipeline integrates state-of-the-art deep learning-based cell segmentation models, including Stardist, to predict cell boundaries. Using these predicted boundaries, several bin-to-cell assignment strategies are applied to obtain single-cell transcript estimates. Cell types are subsequently inferred using either CellAssign (5), a probabilistic method; CellTypist (6), a logistic regression model; or Sargent (7), a scoring-based approach. With the predicted cell types and their locations, downstream analysis can be conducted on the output AnnData objects using Squidpy (8) to identify patterns of cell type co-occurrence, enrichment, and distribution around tissue landmarks.

## Materials and Methods

### Cell Segmentation

Fig. 1 presents the key steps involved in the proposed processing and analysis pipeline. Cells in the full resolution tissue image are first segmented using Stardist, one of the most widely used UNet-based cell segmentation methods that has been shown to accurately segment images even when cells are tightly packed (4).

**Fig. 1.**
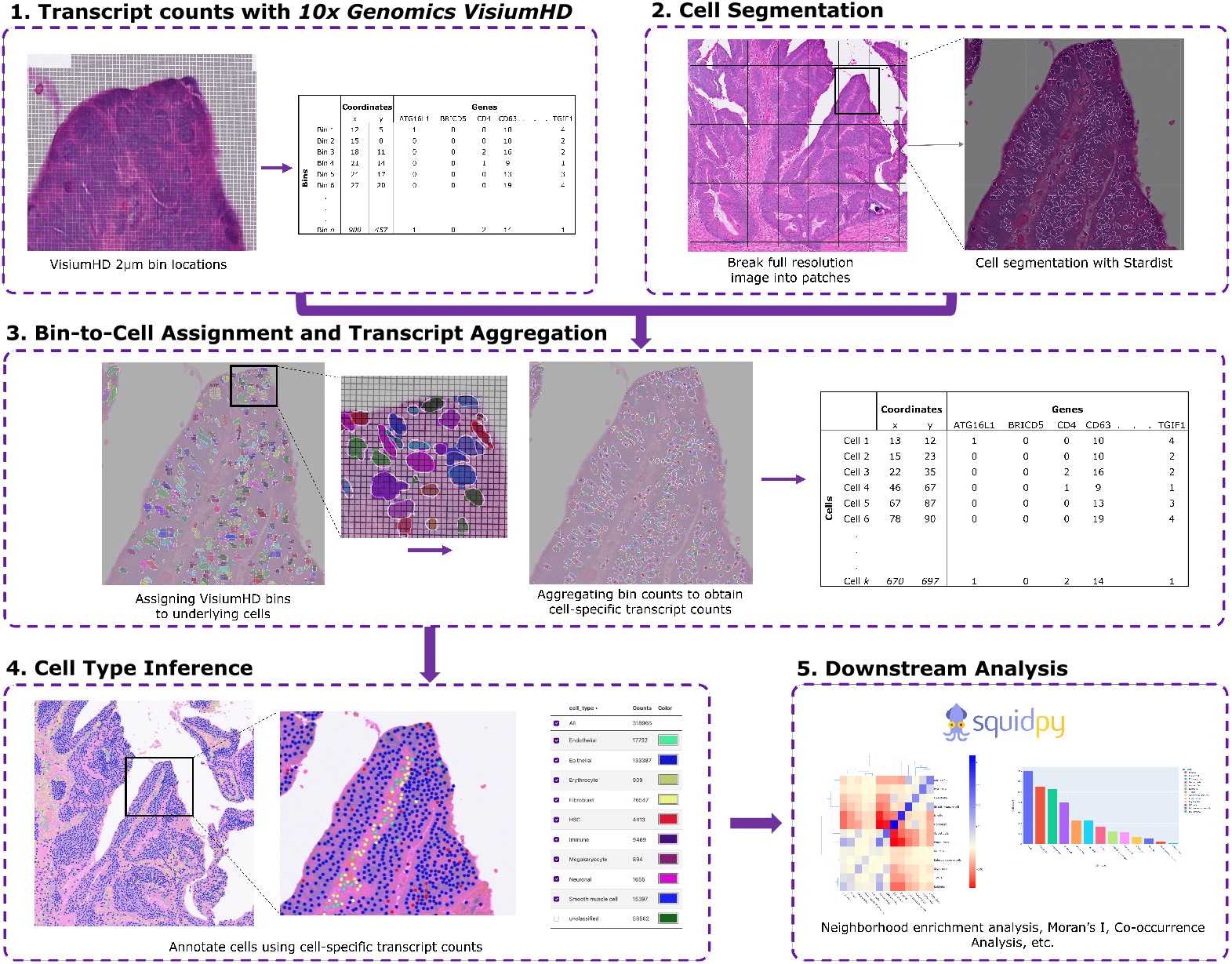
Processing pipeline. (1) Bin-by-gene matrix from 10X Genomics Visium HD provides mRNA transcript counts for each 2μm x 2μm bin. (2) Segmenting the full resolution tissue image using Startdist. (3) Visium HD bins that overlap with a cell are aggregated via a summing operation. This leads to a 2D cell-by-gene matrix. (4) Sargent, CellAssign, or CellTypist are used to translate the cell-wise transcript counts to a respective cell label. (5) Cell labels and their spatial locations within the tissue are wrapped in AnnData objects, which is compatible with SquidPy. This enables the application of various spatial statistical analyses, including neighborhood enrichment, Moran’s I, and co-occurrence analyses. Spatial distribution of the cells can be visualized by Tissuumaps (9) digital pathology viewer.

### Bin-to-Cell Assignment

To obtain the total transcript counts for each cell in the tissue, Visium HD bins that intersect with the inferred cell boundary are aggregated. This produces a 2D cell-by-gene matrix, similar to a single-cell RNA sequencing (scRNA-Seq) experiment with an additional cell ID column that can be mapped to obtain the reading location on the image.

The process begins by representing the Visium HD bins and the predicted cell outlines as Shapely (10) polygons for efficient geometric operations such as computing intersection area and spatial relationships. A spatial join operation is run to identify the spatial relationship between the Visium HD bins and the cell outlines that they intersect. Visium HD bins that do not geometrically intersect with any cell outlines are removed from further analysis. To address bins overlapping multiple cells, we include in ENACT four different transcript imputation methods: a naive method akin to Bin2cell, and three weight-based strategies. Supplementary Section 1 provides details on the implementation of all four bin-to-cell assignment strategies.

### Cell Type Annotation

Several methods are available for cell type annotation using the obtained single cell expression profiles. We use one of Sargent (SignAtuRe-GEne cell aNnoTation) (7), CellAssign (5), or CellTypist (6) to annotate cell types. Sargent is a score-based, single-cell inference algorithm that identifies the cell types of origin based on cell-type specific gene markers. This fits very well for Visium HD data since transcript counts are often low compared to traditional scRNA-Seq experiments, making score-based methods more fitting. CellAssign infers the cell type by computing a probabilistic assignment relative to a cell-type specific gene markers. Lastly, CellTypist is a machine learning-based tool that utilizes classifiers pre-trained on an extensive reference dataset of annotated cells to rapidly assign cell types. CellTypist does not require users to provide gene markers, as they are already defined in the pre-trained models. For Sargent and CellAssign, markers can be obtained from a variety of sources including open-source databases such as PanglaoDB (11) or Cell-Marker (12) .

### Downstream Analysis

The cell labels and their respective locations in the tissue are wrapped in AnnData objects for easy import into open source libraries such as SquidPy. This can be used to run several spatial statistical tests such as neighborhood enrichment analysis, Moran’s I analysis, co-occurrence analysis, etc.

## Evaluation Datasets

To directly evaluate the accuracy of transcript assignment, two synthetic Visium HD-like datasets are constructed from Xenium (13) and sequential fluorescence in situ hybridization (seqFISH+) (14). In addition, the end-to-end pipeline is evaluated against pathologist annotations in the form of anatomical landmarks and manually annotated cell type labels. Details on the synthetic dataset generation and pathologist annotations are provided in Supplementary Sections 2.1 and 2.2, respectively.

## Results

### Evaluating Bin-to-Cell Assignment Methods

While all four possible bin-to-cell assignment methods are available for users, we first evaluate them on two synthetic datasets to provide guidance on their performance.

Results for the Xenium-based synthetic dataset with whole cell boundaries are shown in Table ST3 and Fig. S2. The ‘Weight-By-Area’ method achieves the highest F1 score among the four approaches when evaluating the entire cell boundary. In contrast, the naive method achieves the highest precision as it only considers the unique bins, omitting all the bins shared with multiple cells. However, this approach results in significant information loss, as 25% of the bins overlap with more than one cell, leading to a low recall.

When focusing solely on cell nuclei, we observe a smaller difference in performance between the ‘Naive’ and weighted methods (Fig. S3). This is likely due to the smaller number of bins overlapping with multiple nuclei, limiting the benefits of the weighted methods over the ‘Naive’ method. Additionally, all methods show lower precision on cell nuclei compared to whole cells, which can be attributed to a higher number of non-overlapping bins located at the boundaries of the nuclei, where some transcripts belong to the cell body rather than to the nucleus.

Similarly, for the seqFISH+ synthetic dataset, all four methods demonstrate high precision (close to 1) and recall (close to 0.99, Fig. S4). This performance can likely be attributed to the relatively large size and sparseness of the NIH-3T3 cells, which generally intersect with 200 to 700 bins. Indeed, only about 5% of bins overlapped more than one cell for this dataset. Given the shorter computation time for the ‘Naive’ method, it is recommended in such cases. Table ST4 and Fig. S5 present the run times for the different bin-to-cell assignment methods explored.

### Evaluating Cell-Type Annotation

#### Evaluating Cell-Types in Pathologist Annotations

To evaluate the impact of different bin-to-cell assignment methods on cell type annotation, we compare the annotations generated by our pipeline, employing the four distinct bin-to-cell assignment strategies, against expert-labeled classifications for 20,991 cells. Tables ST7 and ST8 present gene markers used by Sargent and CellAssign to annotate the cell types. This list is curated by a combination of expert provided gene markers and PanglaoDB where the top 10 genes by sensitivity and specificity in humans are selected. To align the more granular cell labels from CellAssign, CellTypist, and Sargent with the broader, pathologist-provided cell labels (Epithelial, Stromal, and Immune cells), the cell types are relabelled according to the mapping defined in Table ST5.

Table ST6 and Fig. S6 summarize the experiments we perform to test the four bin-to-cell assignment methods with the three cell-type annotation algorithms. The performance of the pipeline is assessed as a multi-class classifier, using accuracy, precision, recall, and F1-score as evaluation metrics for the predicted cell types. Table ST6 shows that the weighted methods consistently outperform ‘Naive’ approache across all three cell annotation methods, yielding higher recall, precision, and F1-score values suggesting that a more accurate bin-to-cell assignment method can improve cell annotation. Among the several cell annotation methods, Sargent exhibits the highest performance for annotating cells, followed by CellTypist, with CellAssign ranking last. Sargent achieves the best results when paired with the Weighted-by-Area approach, yielding an accuracy of 0.708 and a weighted F1-score of 0.758.

Fig. S7 presents the confusion matrices for all twelve experiments, highlighting distinct patterns in cell misclassification across methods. CellAssign fails to identify all immune cells, misclassifying them as epithelial cells. Similarly, CellTypist confuses a significant portion of immune cells being incorrectly labeled as epithelial. In contrast, Sargent shows the best performance, significantly reducing misclassifications and accurately differentiating immune cells from epithelial cells.

Errors in the predicted cell types may stem from the cell-specific gene markers used, as these directly influence classification outcomes. Additionally, the manual annotation process, being inherently labor-intensive, may result in mislabeled cells, contributing to the observed inaccuracies.

It is also noteworthy that all methods tend to assign cells as ‘no label’ when the model is uncertain about their classification. This feature could be beneficial in scenarios where researchers aim to identify novel cell types or when the cell types present in a tissue are not well-defined, offering flexibility and the potential to discover new insights.

#### Evaluating Cell-Types in Anatomical Landmarks

To assess the high-level performance of the proposed pipeline, we analyze the assigned cell types within each anatomical landmark labelled by the pathologist.The ‘Weighted-by-Area’ is used for bin-to-cell assignment and Sargent is used for cell type annotation to achieve the highest performance.

For the Human Colorectal Cancer sample, the top predicted cell types in the ‘muscular’ landmark are Smooth Muscle cells, B cells, and Fibroblasts. In the ‘normal epithelial’ region, the predominant cell types identified are Goblet cells, B cells, and Crypt cells. Within the ‘tumoral epithelium’ region, Enterocytes, Goblet cells, and B cells are most prevalent. Finally, the ‘stromal’ regions are characterized primarily by Fibroblasts, B cells, and Endothelial cells.

In the Mouse Small Intestine sample, the gene markers specified in Table ST8 are used by Sargent for cell type annotation.

Smooth muscle cells and Paneth cells are the dominant cell types in the ‘muscular’ landmark. In the ‘normal epithelium’ region, Paneth cells, Goblet cells, and B cells are most frequently observed. The ‘lymphoid’ region is predominantly associated with B cells.

Overall, these results demonstrate significant agreement between the predicted cell types and the associated anatomical landmarks, aligning with established biological knowledge of these tissues (15–17). A limitation of this evaluation approach is that the pathologist annotations obtained might be heterogeneous, for example the regions annotated as ‘epithelial’ can in fact contain smaller stromal or immune aggregate regions that could lead to other non-Epithelial specific cell types being associated with the ‘epithelial’ regions.

## Discussion

ENACT addresses the challenge of spatially resolving cell types in Visium HD data by integrating cell segmentation with high-resolution gene counts; being the first end-to-end pipeline for Visium HD. When evaluated on synthetic data, we find that the weighted methods had better overall precision and recall compared to the ‘Naive’ method. More specifically, we find that the ‘Weighted-by-Area’ which depends on the overlap area for transcript assignment combined with the Sargent cell annotation method performs best. The outputs from the workflow could be used to compute various spatial statistical properties aiding researchers in their ST workflows. The method can also be used to generate large scale datasets combining image and gene data without requiring any manual annotation. This will aid in the training of powerful vision-based cell classifiers on both morphological and gene-based features.

ENACT is available as a python package from our github page: https://github.com/Sanofi-Public/enact-pipeline.

## Acknowledgements

We sincerely appreciate the work of the creators and developers of the various functions, packages, and tools integrated into our workflow and the guidance and feedback from our colleagues at the Precision Medicine and Computational Biology department in Sanofi.

