## Supplementary Material for "ENACT: End-to-End Analysis of Visium High Definition (HD) Data"

**This document provides the supplementary material for ENACT. Code, installation instructions and examples are publicly available on GitHub at https://github.com/Sanofi-Public/enact-pipeline. Experimental data is available at https://zenodo.org/records/13887921.**

#### 1 Bin-to-Cell Assignment Methods

Bin-to-cell assignment begins by omitting the Visium HD bins that do not geometrically intersect any cell outlines. Let $B$ denote the union of the set of remaining bins that completely overlaps a single cell, $B_{\text{unique}}$, and the set of bins that overlap multiple cells, $B_{\text{shared}}$.

$$B = B_{\text{unique}} \cup B_{\text{shared}} \tag{1}$$

The following subsections describe the different strategies developed for bin-to-cell assignment:

##### 1.1 Naive method

This method is similar to Bin2Cell and only uses the $B_{\text{unique}}$ set. Bins that intersect multiple cells ($B_{\text{shared}}$) are omitted.

$$\mathrm{G}_{C_j} = \sum_{i=1}^{N} \alpha_i \times G_{B_i}, \quad \alpha_i = \begin{cases} 1 & \text{if } i \in B_{\text{unique}} \\ 0 & \text{if } i \in B_{\text{shared}} \end{cases} \tag{2}$$

Here $\mathrm{G}_{C_j}$ is the transcript counts assigned to cell $C_j$ and $G_{B_i}$ represents the transcript counts within bin $B_i$. $N$ is the total number of bins that intersect $C_j$.

##### 1.2 Weight-by-Area

The 'naive' method leads to information loss. An alternative approach is to assign bins in $B_{\text{shared}}$ to cells they overlap by weighting the contribution of the bin to the cell based on the area of overlap, $\text{Area}_{C_j \cap B_i}$:

$$\mathrm{G}_{C_j} = \sum_{i=1}^{N} \alpha_i \times G_{B_i}, \quad \alpha_i = \begin{cases} 1 & \text{if } i \in B_{\text{unique}} \\ \frac{\text{Area}_{C_j \cap B_i}}{\text{Area}_{B_i}} & \text{if } i \in B_{\text{shared}} \end{cases} \tag{3}$$

Here, $\text{Area}_{B_i}$ is the total bin area ($\sim 4\ \mu\text{m}^2$).

##### 1.3 Weighted-by-Transcript

While area weighting allows the assignment of all overlapping bins, such an assignment ignores the fact that neighboring cells can be very different. Thus, simply dividing the transcripts based on area may not provide the correct assignment of these transcripts. This method attempts to address that by dividing the transcripts based on the expression of the cells overlapping the bin:

$$
\begin{aligned}
\mathrm{G}_{C_j} &= \sum_{i=1}^{N} \left( \sum_{g=1}^{T} \alpha_{i,g} \times G_{B_i,g} \right), \\
\alpha_{i,g} &= \begin{cases} 1 & \text{if } i \in B_{\text{unique}} \\ \frac{\hat{G}_{C_j,g}}{\sum_{s=1}^{n} \hat{G}_{C_s,g}} & \text{if } i \in B_{\text{shared}} \end{cases}
\end{aligned} \tag{4}
$$

where:

- $T$ is the total number of unique genes,
- $\alpha_{i,g}$ is the weighting for gene $g$ in bin $i$,
- $G_{B_i,g}$ is the count of gene $g$ in bin $B_i$,
- $\hat{G}_{C_j,g}$ is the normalized count of gene $g$ in cell $C_j$, and
- $n$ is the number of cells that overlap with bin $B_i$.

This method incorporates transcript information from surrounding cells to accurately distribute the transcript counts from bins shared with multiple cells. The weighting term, $\alpha_{i,g}$, is applied according to the proportion of normalized transcript counts, $\hat{G}$, in the overlapping cells.

#### 1.4 Weighted-by-Cluster

One problem with the 'Weighted-by-Transcript' method is that if an overlapping bin contains a gene that has not been expressed in any of the other bins of the intersecting cells, the weighting factor $\alpha_{i,g}$ would be 0 for all cells leading to information loss. This may impact the assignment of many genes expressed at low levels which can still play an important role in the process being studied (1). To address this, here $\alpha$ is computed by using the (average) expression of the cell type they belong to. Specifically, we first use the 'Naive' method to obtain individual cell transcript estimates. K-means clustering is used to group cells based on gene expression. For each cluster, the average transcript counts are computed and used for assigning the remaining bins.

$$
\begin{aligned}
\mathrm{G}_{C_j} &= \sum_{i=1}^{N} \left( \sum_{g=1}^{T} \alpha_{i,g} \times G_{B_i,g} \right), \\
\alpha_{i,g} &= \begin{cases} 1 & \text{if } i \in B_{\text{unique}} \\ \frac{\hat{G}_{\mathcal{K}(C_j),g}}{\sum_{s=1}^{n} \hat{G}_{\mathcal{K}(C_s),g}} & \text{if } i \in B_{\text{shared}} \end{cases} \\
\mathcal{K}(C) &= \text{K-means}(C)
\end{aligned} \tag{5}
$$

where $\mathcal{K}(C)$ is the cluster that cell $C$ is assigned to.

### 2 Evaluation Datasets

#### 2.1 Datasets for Evaluating Bin-to-Cell Assignment Methods

To directly evaluate the accuracy of transcript assignment, two synthetic Visium HD-like datasets are constructed. The first dataset is constructed from Xenium datasets provided by 10x Genomics, profiling FFPE Human Colorectal Cancer (Xenium demo data[1]). This dataset includes 386,695 segmented cells/nuclei and 545 distinct genes. As Xenium is an imaging-based technology that provides pinpoint locations of transcripts within the tissue, it serves as a good baseline for assessing whether our bin-to-cell methods accurately map transcript locations.

The second dataset is from sequential fluorescence in situ hybridization (seqFISH+). The seqFISH+ dataset (2) contains spatial mRNA data for 10,000 genes, offering gene coverage that is more akin to Visium HD. Like Xenium, this is also an imaging method making it easier to assign specific transcripts to cells. The dataset contains data from 103 mouse embryonic fibroblast (NIH-3T3) where cells were manually segmented[2].

To generate synthetic datasets from these two datasets we artificially assign a 2µm x 2µm grid to the image and summed up the transcript abundance in each bin. Additionally, since our segmentation model, Stardist, focuses on nuclei segmentation and the Xenium dataset provides information on both nuclei and whole cell boundaries, we assess the assignment methods by comparing their performance in assigning transcripts to both nuclei and whole cell boundaries.

#### 2.2 Datasets for Evaluating Cell-Type Annotation

To validate the end-to-end pipeline, two publicly available FFPE Visium HD samples are analyzed: human colorectal cancer[3] and mouse small intestine[4]. Pathologist annotations are obtained in the form of anatomical landmarks and manually annotated cell type labels as shown in Fig. S1. Table ST1 presents the breakdown of the different anatomical landmarks labelled by the pathologists. Table ST2 presents the breakdown of the cell types manually labelled by the pathologist. In total 20,991 manually curated cell labels were obtained from four tissue patches in the Human Cancer samples.

| Sample Type | Tissue Landmarks |
|---|---|
| Human colorectal cancer[5] | Muscular, Normal Epithelium, Tumoral Epithelium, Stroma |
| Mouse small intestine[6] | Lymphoid Tissue, Muscular, Normal Epithelium |

**Table 1.** List of sample types and their corresponding tissue anatomical landmarks.

| Cell Type | Number of Annotated Cells |
|---|---|
| Epithelial cells | 12072 |
| Stromal cells | 6171 |
| Immune cells | 2748 |

**Table 2.** List of cell types and their corresponding count in the evaluation dataset

[1] https://www.10xgenomics.com/datasets/ffpe-human-colorectal-cancer-data-with-human-immuno-oncology-profiling-panel-and-custom-add-on-1-standard
[2] https://zenodo.org/records/2669683#.Xqi1w5NKg6g
[3] https://www.10xgenomics.com/datasets/visium-hd-cytassist-gene-expression-libraries-of-human-crc
[4] https://www.10xgenomics.com/datasets/visium-hd-cytassist-gene-expression-libraries-of-mouse-intestine

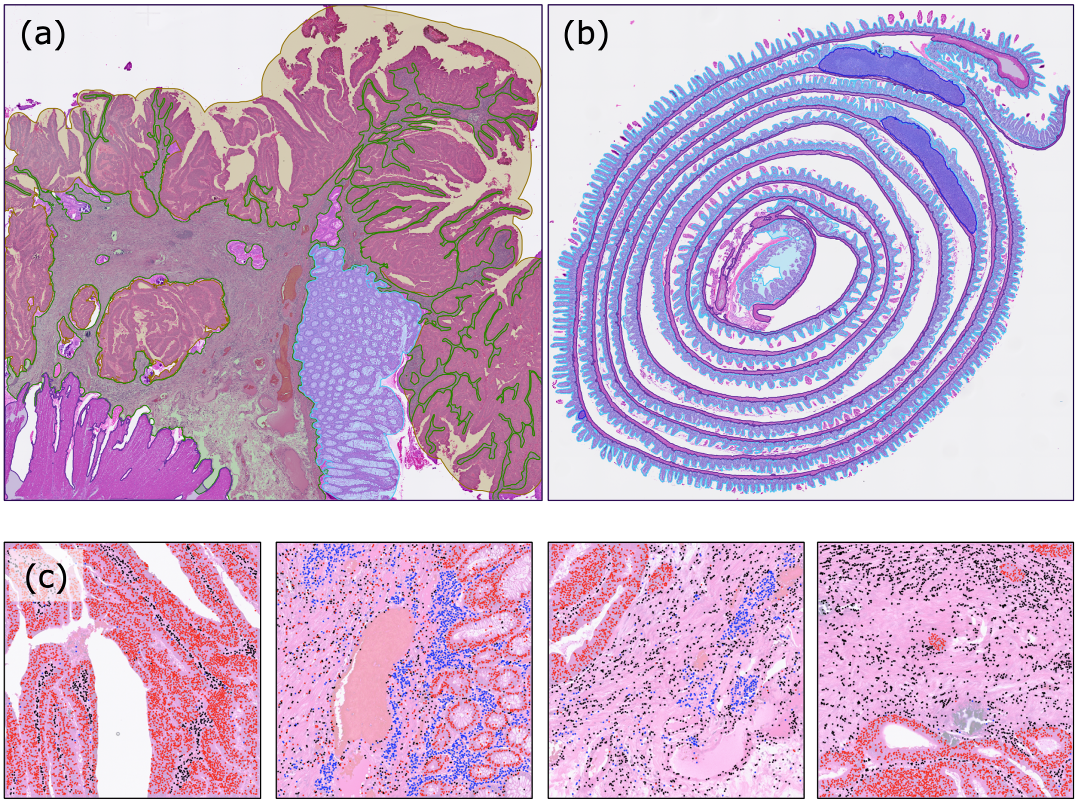


**Fig. 1.** (a) Human Colorectal cancer sample with pathologist-annotated anatomical landmarks for Tumoral Epithelium (yellow), Normal Epithelium (blue), Muscular (purple), and Stroma (green) areas. (b) Mouse small intestine sample with pathologist-annotated anatomical landmarks for Lymphoid Tissue (purple), Muscular (pink), Normal Epithelium (blue) areas. (c) Patches from Human Colorectal cancer sample with individually annotated cell labels for Epithelial cells (red), Immune cells (blue), and Stromal cells (black).

### 3 Results

#### 3.1 Evaluating Bin-to-Cell Assignment Methods

This section describes the performance of the different bin-to-cell assignment methods on the Xenium- and (SeqFISH+)- based synthetic datasets. For the Xenium-based dataset, both the whole cell and nuclei boundaries are used for bin-to-cell assignment. For the SeqFISH+ dataset, only whole cell boundaries are considered since the nuclei boundaries are not provided.

##### 3.1.1 Performance on Xenium-Based Synthetic Dataset

| Method | Precision Avg. | Recall Avg. | F1 Avg. |
|---|---|---|---|
| Naive | **0.98** | 0.48 | 0.65 |
| Weighted-by-Area | 0.90 | **0.87** | **0.88** |
| Weighted-by-Transcript | 0.82 | 0.70 | 0.76 |
| Weighted-by-Cluster | 0.812 | 0.83 | 0.82 |

**Table 3.** Evaluation of the different bin-to-cell assignment methods on the Xenium-based synthetic dataset using the whole cell boundaries.

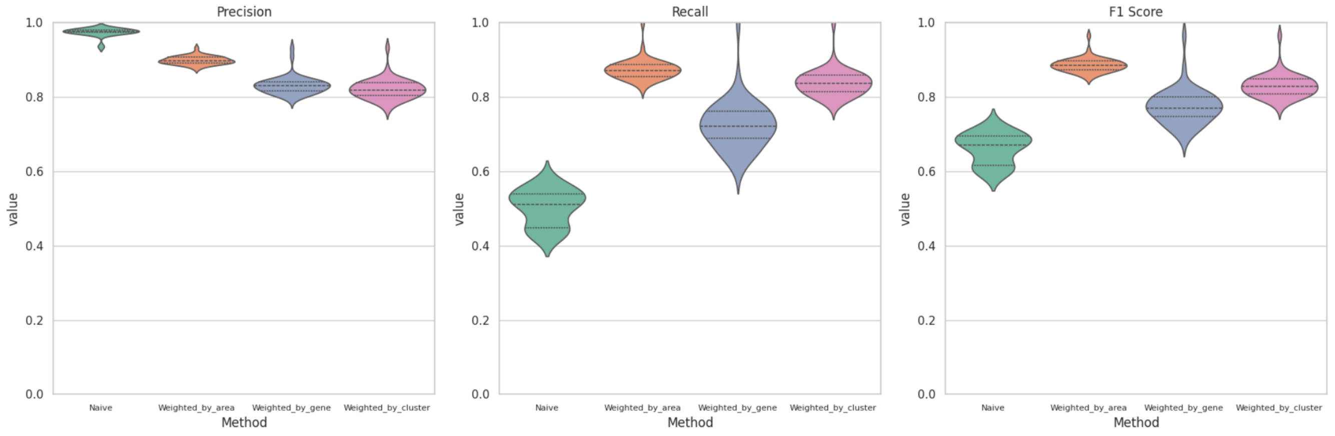


**Fig. 2.** Violin plots of the precision, recall, and F1 score of the four bin-to-cell assignment methods evaluated on the Xenium-based synthetic dataset using the whole cell boundaries.

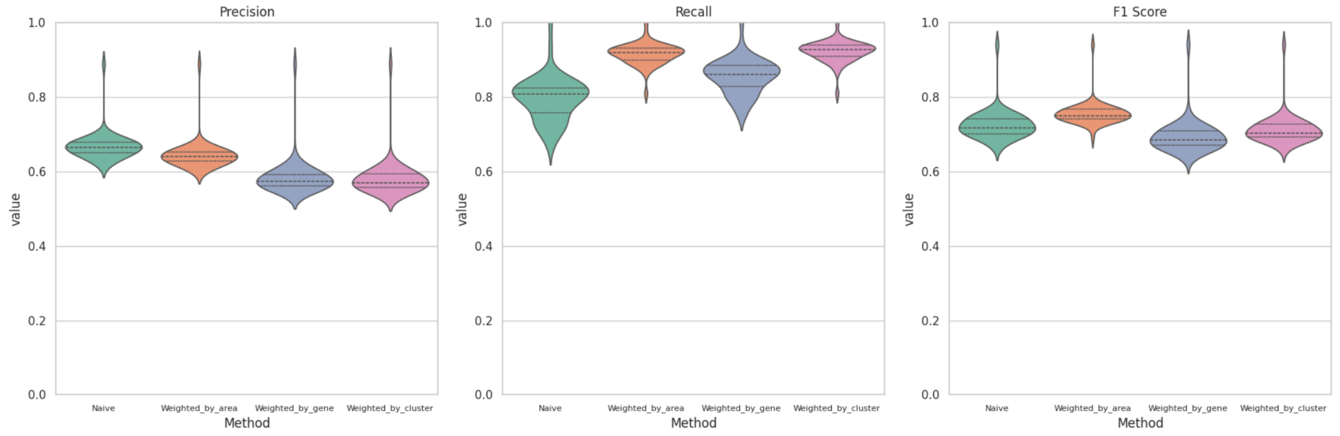


**Fig. 3.** Violin plots of the precision, recall, and F1 score of the four bin-to-cell assignment methods evaluated on the Xenium-based synthetic dataset using the nuceli boundaries.

##### 3.1.2 Performance on (seqFISH+)-Based Synthetic Dataset

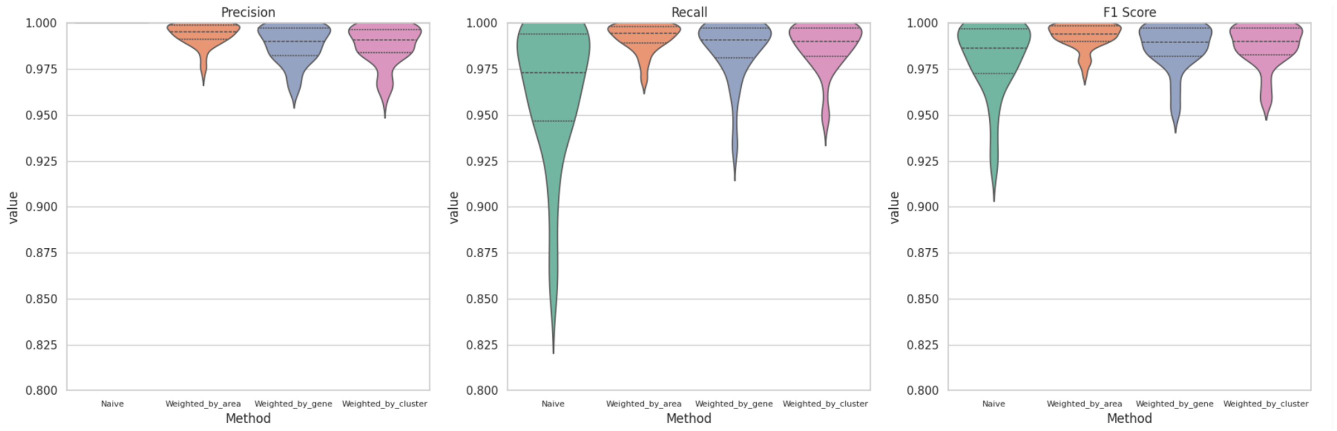


**Fig. 4.** Violin plots of the precision, recall, and F score of the four bin-to-cell assignment methods evaluated on the (SeqFISH+)-based synthetic dataset using the whole cell boundaries.

#### 3.2 Bin-to-Cell Assignment Method Running Time Analysis

The run time for the four bin-to-cell assignment methods is computed by measuring time needed to perform bin-to-cell assignment on patches of size 4000 x 4000 pixels from the Human colorectal cancer public sample [7]. In total, 42 patches are analyzed, each containing around 10,000 cells. Here, only the top-1000 most highly variable genes are used used for the analysis.

| | Running Time Avg. (Second) |
|---|---|
| **Naive** | 7.01 |
| **Weighted-by-Area** | 15.2 |
| **Weighted-by-Transcript** | 61.3 |
| **Weighted-by-Cluster** | 78.0 |

**Table 4.** This table summarizes the execution time (in seconds) for each bin-to-cell assignment method when processing a 4000 x 4000 pixel image patch with around 10000 cells.

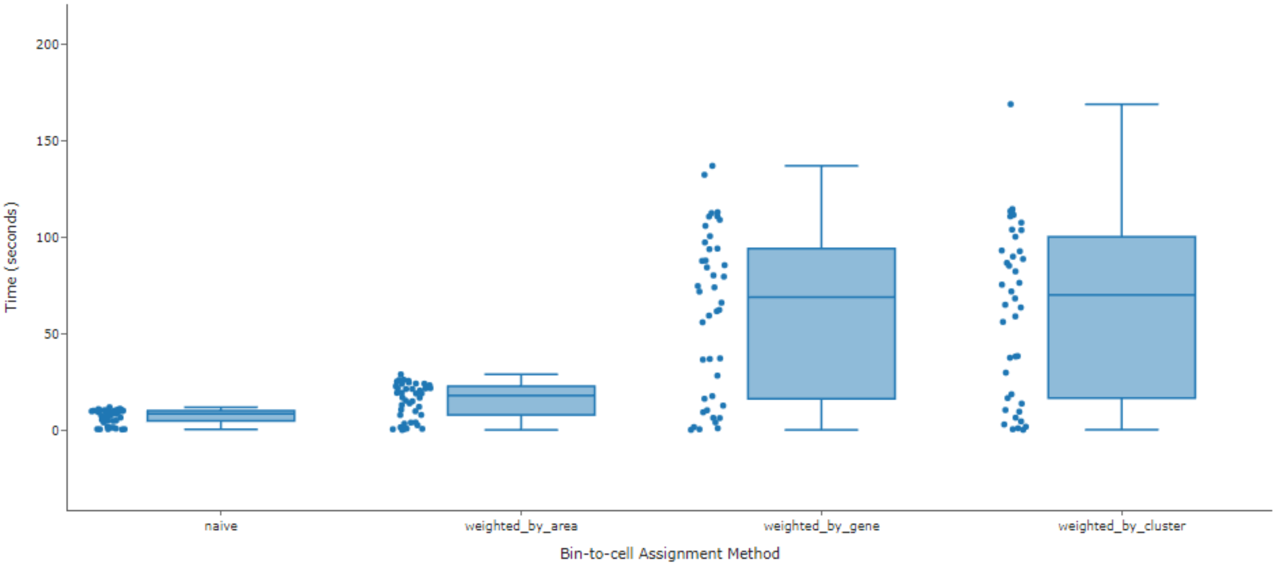


**Fig. 5.** Execution time (in seconds) for each bin-to-cell assignment method when processing a 4000 x 4000 pixel image patch with around 10000 cells. Here, each blue dot represents the run time for a given patch.

[7] https://www.10xgenomics.com/datasets/visium-hd-cytassist-gene-expression-libraries-of-human-crc

##### 3.3 Comparing Predicted Cell Types to Pathologist Annotations

The predictions from ENACT are compared against pathologist-curated cell labels from the Human colorectal cancer public sample. Here, twelve experiments are conducted to compare the effect of the bin-to-cell assignment method and cell annotation algorithm on the pipeline performance. The top-1000 most highly variable genes in addition to the gene markers defined in Table ST7 are used for bin-to-cell assignment and cell annotation. Table ST5 describes the mapping between the granular cell labels predicted by Sargent, CellAssign, and CellTypost and the pathologist-provided cell label. Table ST6 provides the summary of the evaluation in terms of accuracy, precision, recall, and F1-score.

<table>
<tr><th>Cell Annotation Method</th><th>Granular label</th><th>Pathologist label</th></tr>
<tr><td rowspan="3">Sargent, CellAssign</td><td>Epithelial, Enterocytes, Goblet cells, Enteroendocrine cells, Crypt cells</td><td>epithelial cells</td></tr>
<tr><td>B cells, T cells, NK cells</td><td>immune cells</td></tr>
<tr><td>Endothelial, Fibroblast, Smooth muscle cell</td><td>stromal cells</td></tr>
<tr><td rowspan="3">CellTypist</td><td>CMS1, Mature Enterocytes type 2, Mature Enterocytes type 1, CMS3, CMS4, Stem-like/TA, CMS2, Goblet cells</td><td>epithelial cells</td></tr>
<tr><td>cDC, Regulatory T cells, Neutrophils, Gamma delta T cells, Macrophages, Eosinophils, CD8+ T cells, CD4+ T cells, IgG+ Plasma, Pro-inflammatory, T helper 17 cells, CD19+CD20+ B, Mast cells, NK cells, T follicular helper cells, IgA+ Plasma</td><td>immune cells</td></tr>
<tr><td>Proliferative ECs, Lymphatic ECs, Stromal 2, Stromal 3, Stalk-like ECs, Tip-like ECs, Smooth muscle cells, Stromal 1, Myofibroblasts, Pericytes, Enteric glial cells</td><td>stromal cells</td></tr>
</table>

**Table 5.** Look up table mapping the granular cell types to the broader pathologist-provided cell type label.

| Cell Annotation Method | Bin-to-Cell Method | Accuracy | Precision | Recall | F-Score |
|---|---|---|---|---|---|
| CellAssign | Naive | 0.581 | 0.570 | 0.581 | 0.467 |
| | Weight-by-Area | 0.581 | 0.569 | 0.581 | 0.468 |
| | Weight-by-Transcript | 0.582 | 0.611 | 0.582 | 0.471 |
| | Weight-by-Cluster | 0.583 | 0.570 | 0.583 | 0.472 |
| CellTypist | Naive | 0.579 | 0.690 | 0.579 | 0.611 |
| | Weight-by-Area | 0.581 | 0.691 | 0.581 | 0.612 |
| | Weight-by-Transcript | 0.580 | 0.690 | 0.580 | 0.611 |
| | Weight-by-Cluster | 0.582 | 0.692 | 0.582 | 0.614 |
| Sargent | Naive | 0.684 | 0.836 | 0.684 | 0.740 |
| | Weight-by-Area | **0.708** | 0.840 | **0.708** | **0.758** |
| | Weight-by-Transcript | 0.688 | 0.838 | 0.688 | 0.744 |
| | Weight-by-Cluster | 0.703 | **0.841** | 0.703 | 0.754 |

**Table 6.** Performance comparison of pipeline with different Bin-to-Cell methods and cell type inference algorithms using the Visium HD Dataset.

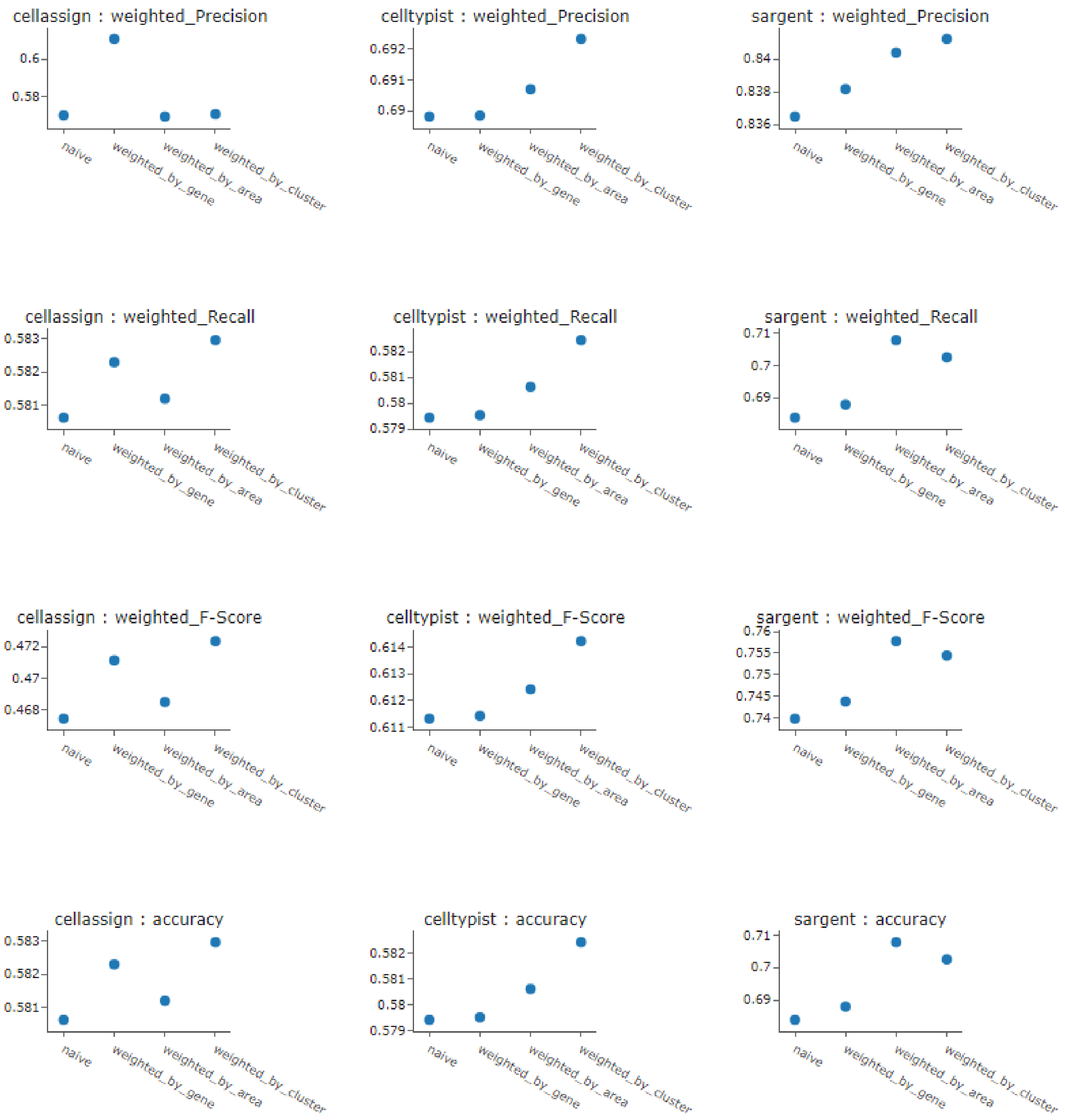


**Fig. 6.** Precision, recall, F score, and accuracy for the twelve experiments run combining the three cell annotation methods: CellAssign, CellTypist, and Sargent, and the four bin-to-cell assignment methods: Naive, Weighted-by-Area (weighted_by_area), Weighted-by-Transcript (weighted_by_gene), and Weighted-by-Cluster (weighted_by_cluster)

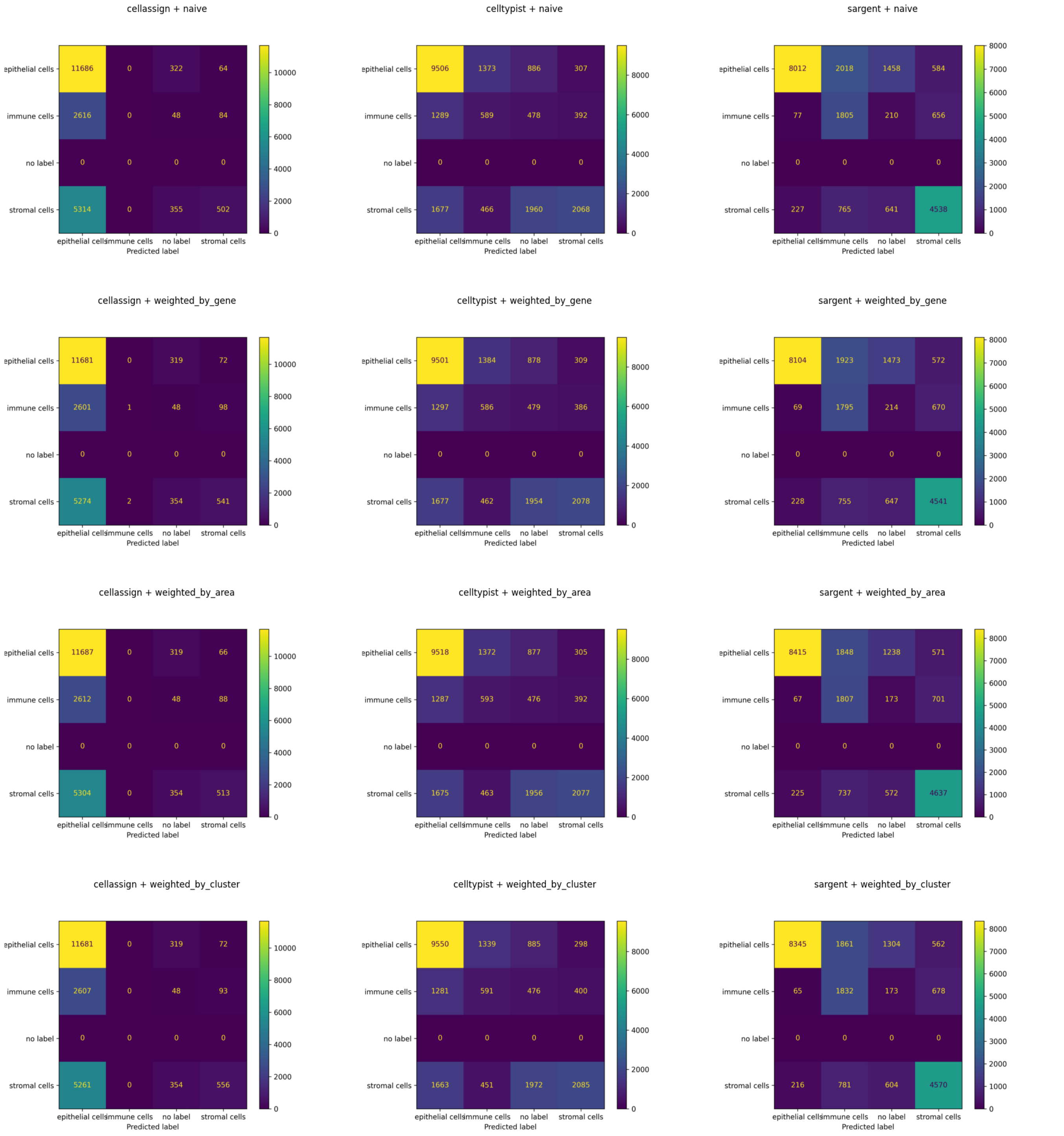


**Fig. 7.** Confusion matrices for the twelve experiments run combining the three cell annotation methods and four bin-to-cell assignment methods: Naive, Weighted-by-Area (weighted_by_area), Weighted-by-Transcript (weighted_by_gene), and Weighted-by-Cluster (weighted_by_cluster)

#### 3.4 Evaluating Cell-Types in Anatomical Landmarks

This section describes a higher-level analysis of the performance of ENACT compared to the analysis provided in Supplementary Section 3.3. Here, the cell-type annotations predicted by ENACT are analyzed in each of the pathologist-annotated anatomical landmarks defined in Table ST1. Only the cells that are located within each anatomical landmark are considered. Tables ST7 and ST8 define the gene markers used for this analysis. Figures S8 and S10 show the spatial organization of the predicted cell type on TissUUmaps histopathology web-viewer. (3).

##### 3.4.1 Human Colorectal Cancer

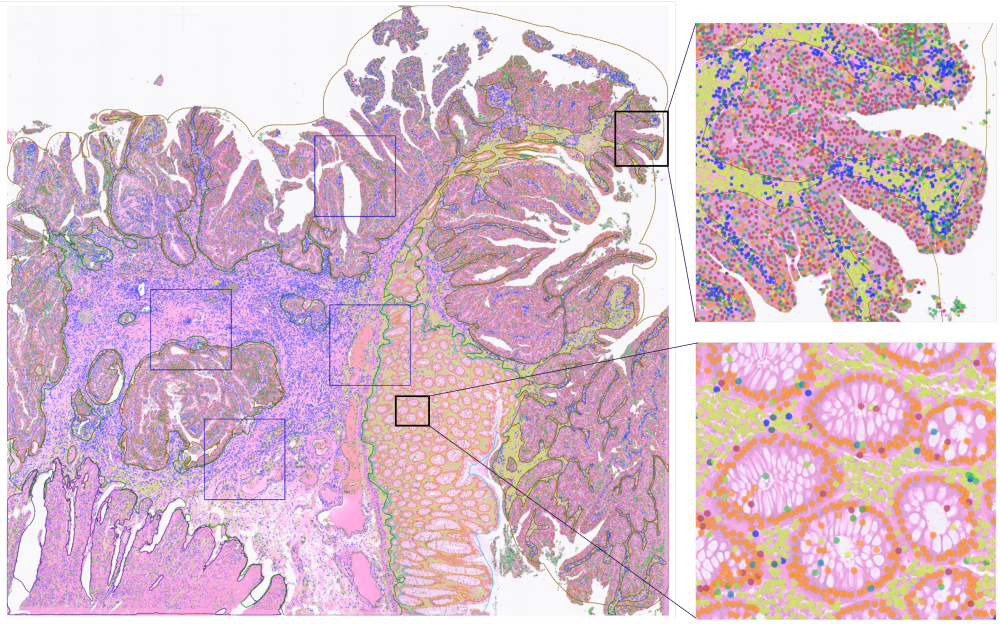

| cell_type | Counts ▾ | Color |
|---|---|---|
| All | 318965 | |
| B cells | 69854 | |
| Fibroblast | 54931 | |
| Enterocytes | 52681 | |
| Goblet cells | 40117 | |
| unclassified | 22887 | |
| Epithelial | 22755 | |
| T cells | 16858 | |
| Smooth muscle cell | 12123 | |
| Endothelial | 11262 | |
| Crypt cells | 6933 | |
| NK cells | 5195 | |
| Enteroendocrine cells | 2361 | |
| Erythrocyte | 1008 | |

**Fig. 8.** Spatial distribution of cell types present in the Human Colorectal sample. Each dot represents the centroid of the cell.

| Epithelial | Enterocytes | Goblet cells | Entero-endocrine cells | Crypt cells | Endothelial | Fibroblast | Smooth muscle cells | B cells | T cells | NK cells |
|---|---|---|---|---|---|---|---|---|---|---|
| CDH1 | CD55 | MANF | NUCB2 | HOPX | PECAM1 | COL1A1 | BGN | CD74 | JUNB | S100A4 |
| EPCAM | ELF3 | KRT7 | FABP5 | SLC12A2 | CD34 | COL3A1 | MYL9 | HMGA1 | S100A4 | IL32 |
| CLDN1 | PLIN2 | AQP3 | CPE | MSI1 | KDR | COL5A2 | MYLK | CD52 | CD52 | CXCR4 |
| CD2 | GSTM3 | AGR2 | ALCAM | SMOC2 | CDH5 | PDGFRA | FHL2 | PTPRC | PFN1P1 | FHL2 |
| | KLF5 | BACE2 | GCG | OLFM4 | PROM1 | ACTA2 | ITGA1 | HLA-DRA | CD81 | IL2RG |
| | CBR1 | TFF3 | SST | ASCL2 | PDPN | TCF21 | ACTA2 | CD24 | EEF1B2P3 | CD69 |
| | APOA1 | PHGR1 | CHGB | PROM1 | TEK | FN | EHD2 | CXCR4 | CXCR4 | CD7 |
| | CA1 | MUC4 | IAPP | BMI1 | FLT1 | | OGN | SPCS3 | CREM | NKG7 |
| | PDHA1 | MUC13 | CHGA | EPHB2 | VCAM1 | | SNCG | LTB | IL32 | CD2 |
| | EHF | GUCA2A | ENPP2 | LRIG1 | PTPRC | | FABP4 | IGKC | TGIF1 | HOPX |
| | | | | | VWF | | | | | |
| | | | | | ENG | | | | | |
| | | | | | MCAM | | | | | |
| | | | | | ICAM1 | | | | | |
| | | | | | FLT4 | | | | | |

**Table 7.** Gene markers used to obtain cell type labels using Sargent for Human Colorectal Cancer sample.

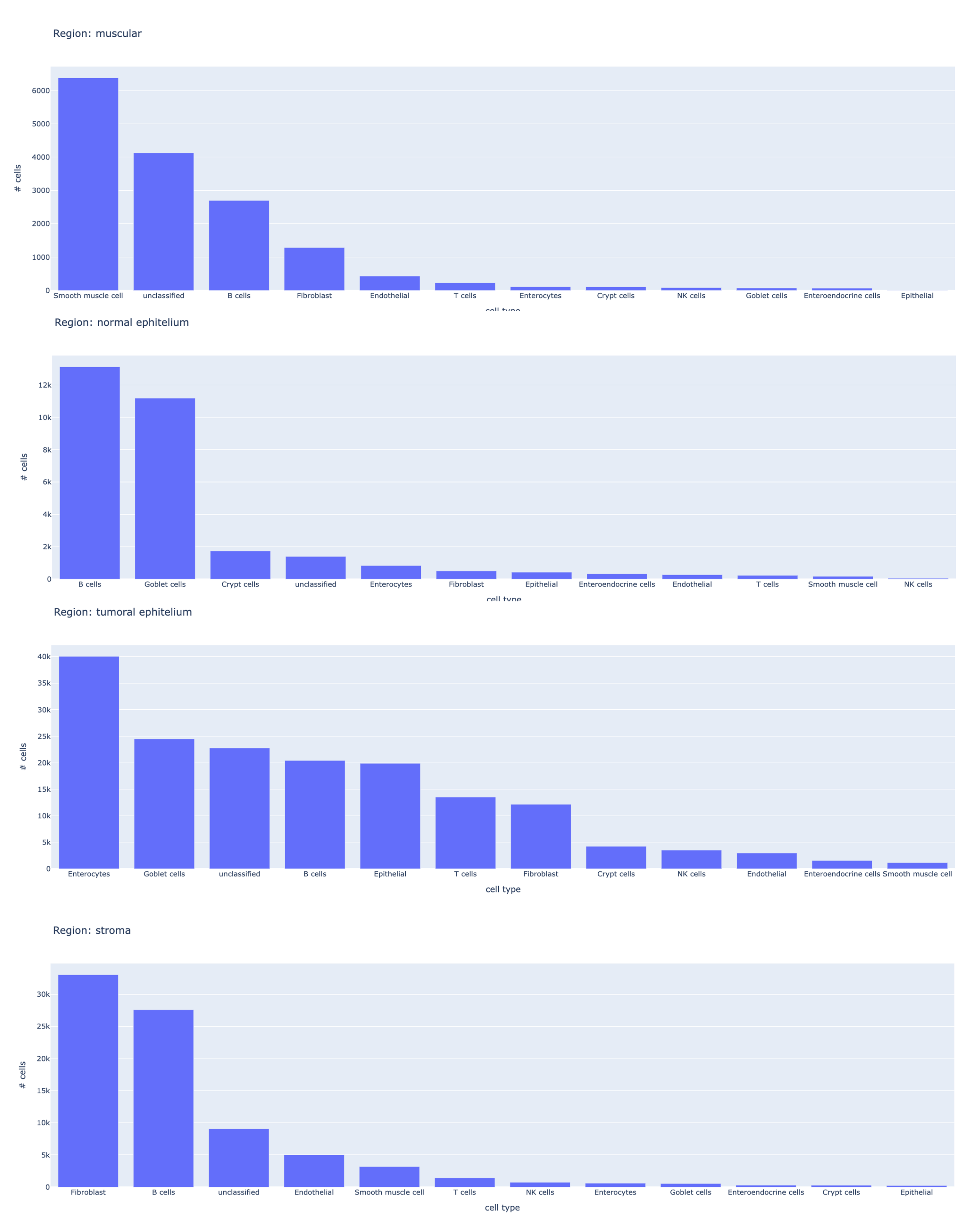


**Fig. 9.** Distribution of the cell types within the four anatomical landmarks annotated in the Human Colorectal sample.

##### 3.4.2 Mouse Small Intestine

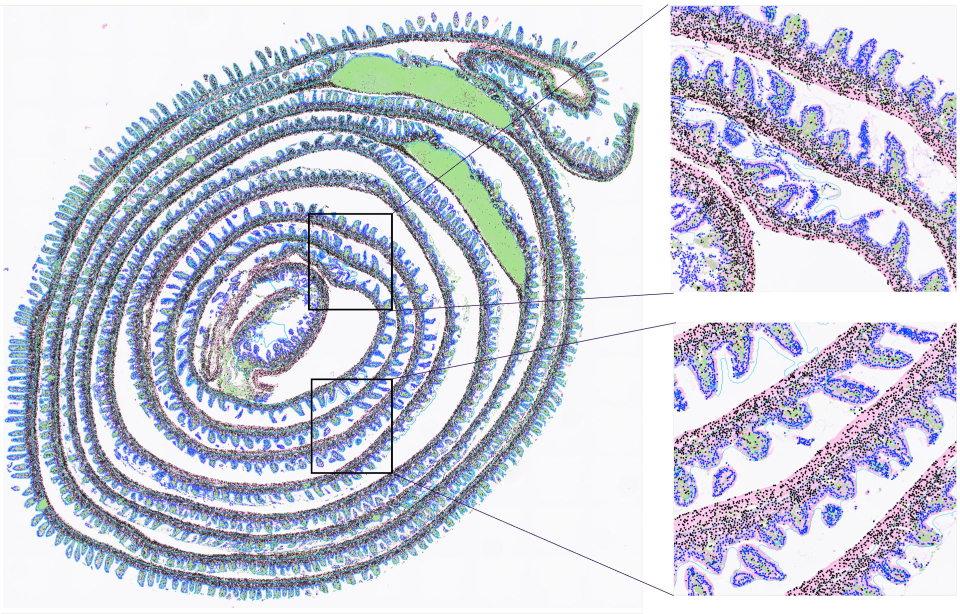

| cell_type ▾ | Counts | Color |
|---|---|---|
| All | 392234 | |
| B cells | 94448 | |
| Crypt cells | 3091 | |
| Enterocytes | 15409 | |
| Enteroendocrine cells | 950 | |
| Goblet cells | 83846 | |
| NK cells | 1978 | |
| Paneth cells | 80831 | |
| Smooth muscle cells | 67796 | |
| T cells | 1546 | |
| unclassified | 42339 | |

**Fig. 10.** Spatial distribution of cell types present in the Mouse Small Intestine sample. Each dot represents the centroid of the cell.

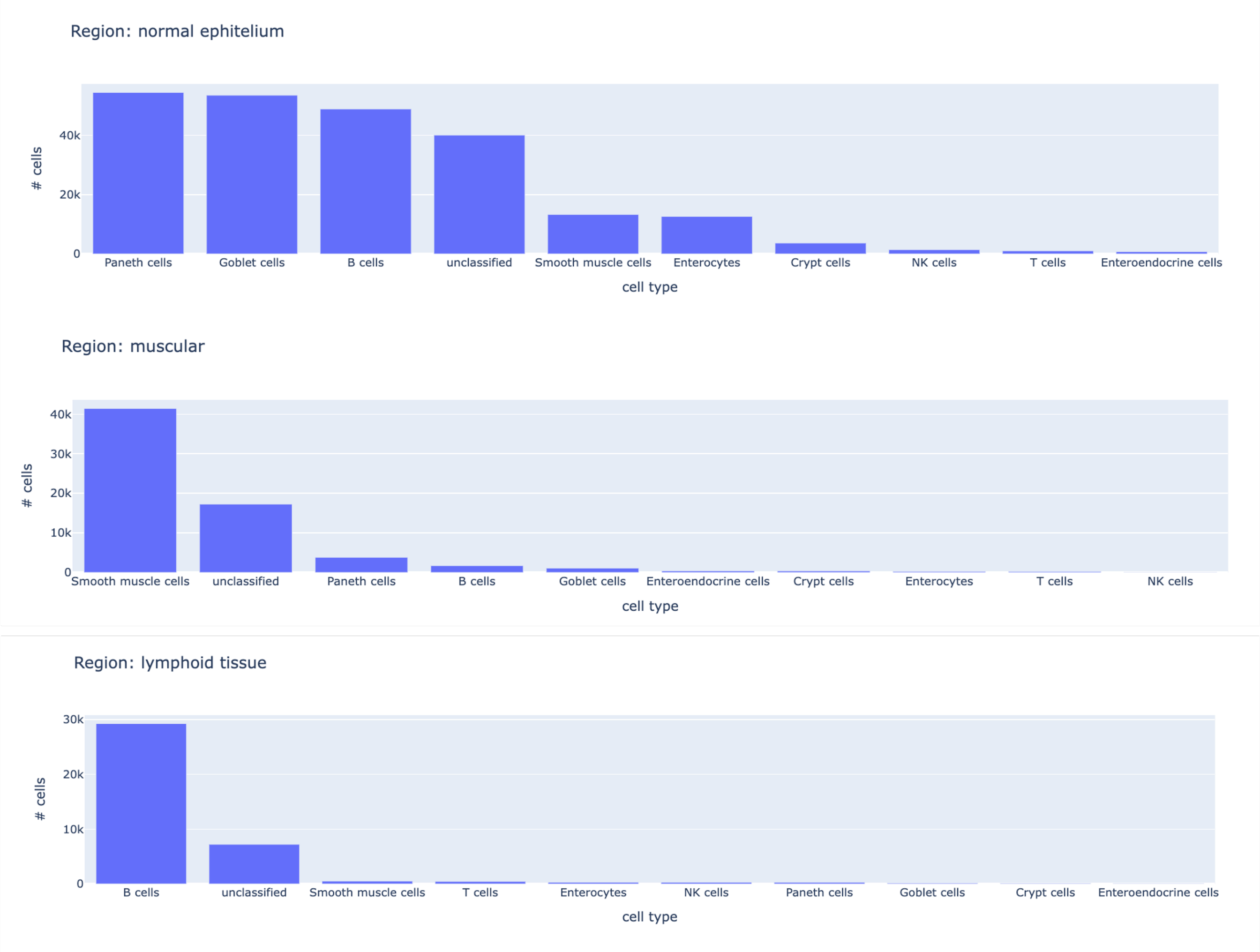


**Fig. 11.** Distribution of the cell types within the four anatomical landmarks annotated in the Mouse Small Intestine sample.

| Enterocytes | Goblet cells | Entero-endocrine cells | Paneth cells | Crypt cells | Smooth muscle cells | B cells | T cells | NK cells |
|---|---|---|---|---|---|---|---|---|
| Cbr1 | Manf | Fabp5 | Gpx2 | Prom1 | Bgn | Cd52 | Cd81 | Ctla2A |
| Plin2 | Krt7 | Cpe | Fabp4 | Hopx | Myl9 | Bcl11A | Junb | Ccl4 |
| Gls | Ccl9 | Enpp2 | Lyz1 | Msi1 | Pcp4L1 | Ebf1 | Cd52 | Cd3G |
| Plin3 | Muc13 | Chgb | Kcnn4 | Olfm4 | Itga1 | Cd74 | Ptprcap | Ccl3 |
| Dab1 | Phgr1 | Alcam | Lgals2 | Kcne3 | Nrp2 | Ptprc | H2-Q7 | Nkg7 |
| Pmepa1 | Cdx2 | Chga | Guca2B | Bmi1 | Mylk | Pold4 | Ccl6 | Lat |
| Acsl5 | Aqp3 | Pax6 | Lgr4 | Axin2 | Ehd2 | Ighm | Bcl2 | Dusp2 |
| Hmox1 | Creb3L1 | Neurod1 | Defa24 | Kcnq1 | Fabp4 | Cd14 | Maff | Itgam |
| Abcg2 | Guca2A | Cck | Il4Ra | Ascl2 | Acta2 | Creld2 | Ccl4 | Fhl2 |
| Cd36 | Klk1 | Isl1 | Guca2A | Lrig1 | Ogn | Fli1 | Ccl3 | Ccl5 |

**Table 8.** Gene markers used to obtain cell type labels using Sargent for Mouse Small Intestine sample.
